# Early-life Temperature Exposure Affects Thyroid Hormone Receptor Signaling and Epigenetic Regulation of the Paraventricular Nucleus in Female Rat Pups

**DOI:** 10.1101/2020.05.01.072488

**Authors:** Samantha C. Lauby, Patrick O. McGowan

## Abstract

Early-life maternal care received has a profound effect on later-life behaviour in adult offspring and previous studies have suggested epigenetic mechanisms (e.g., DNA methylation) are involved. Changes in thyroid hormone receptor signaling might be related to differences in maternal care received and DNA methylation modifications. We investigated the effects of two factors in the maternal environment, temperature exposure (a proxy of maternal contact) and licking-like tactile stimulation, on these processes in week-old female rat pups. We assessed thyroid hormone receptor signaling by measuring circulating triiodothyronine and transcript abundance of thyroid hormone receptors and the thyroid hormone-responsive genes DNA methyltransferase 3a and oxytocin in the paraventricular nucleus of the hypothalamus. DNA methylation of the oxytocin promoter was assessed in relation to changes in thyroid hormone receptor binding. Repeated room temperature exposure was associated with a decrease in thyroid hormone receptor signaling measures relative to nest temperature exposure, while acute room temperature exposure was associated with an increase. Repeated room temperature exposure also increased thyroid hormone receptor binding and DNA methylation at the oxytocin promoter. These findings suggest that repeated room temperature exposure may affect DNA methylation levels as a consequence of alterations in thyroid hormone receptor signaling.

## 1. Background

Maternal care in early life, typically assessed by licking/grooming in rodents, has a profound influence on neurodevelopmental trajectories in offspring and their later-life behaviour [1,2]. Early-life maternal separation, which disrupts maternal-pup contact, or natural variations in maternal care persistently alter transcript abundance of corticotropin releasing factor (*Crf*) [3], arginine vasopressin (*Avp*) [4], and the glucocorticoid receptor [5,6] in the brain. These changes in transcript abundance have been linked to persistent epigenetic modifications of DNA and histones that alter the binding of transcription factors to DNA. In addition, other maternal factors may play a role in later-life offspring phenotypes either by themselves or via interactions with licking/grooming [2,7]. The exposure of pups to lower ambient (room) temperatures as a result of brief disruptions in mother-pup contact has been proposed to be an important component involved in the reduction in stress response following early-life handling [7–9]. Pups have an inefficient thermoregulatory system [10] and are dependent on non-shivering thermogenesis, huddling with siblings, and proximal contact with the rat mother [10–12] to maintain body temperature. Licking-like tactile stimulation has also been shown to reduce the body temperature of rat pups [13].

Variations in both ambient temperature and licking/grooming alter thyroid hormone physiology. Room temperature exposure can induce release of thyroid hormones to evoke non-shivering thermogenesis in brown adipose tissue. In addition, there is some evidence that acute licking-like tactile stimulation can induce conversion of the thyroid hormone thyroxine to triiodothyronine (T3) and lead to downstream DNA demethylation of the glucocorticoid receptor promoter in the hippocampus [14].

Thyroid hormones, especially T3, play important roles in neurodevelopment by regulating gene transcription through thyroid hormone receptor [15]. In the developing mouse brain, transcription of DNA methyltransferase 3a (*Dnmt3a*), an enzyme that catalyzes *de novo* DNA methylation modifications, can occur through the binding of liganded thyroid hormone receptor to thyroid hormone response elements (TREs) in intragenic regions of *Dnmt3a* [16]. Interestingly, mouse pups that receive higher levels of maternal care also show increased DNA methyltransferase 3a transcript abundance in the hippocampus at postnatal day 7 [17]. Thyroid hormone is also a regulator of oxytocin (*Oxt*) [18], a neuropeptide produced in the paraventricular nucleus (PVN) of the hypothalamus and involved with stress attenuation, among other phenotypes. Previous work has demonstrated that thyroid hormone and other receptors can bind to the composite hormone response element (CHRE) in the promoter region of *Oxt to* activate transcription [18–20]. Genetic deletion of this region abolishes transcription, indicating the CHRE is required for proper regulatory control of *Oxt* [21]. A CpG dinucleotide flanks the 5’ end of the CHRE but its potential role in the epigenetic regulation of *Oxt* in the PVN has not been examined. Previous work in humans has investigated DNA methylation in the entire *Oxt* promoter region, including the CHRE [22,23], and the enhancer in the *Oxt/Avp* intergenic region [24]. However, because peripheral tissues were examined in these studies, it is not known whether DNA methylation at this locus is involved in the regulation of *Oxt* transcript abundance in the brain.

In this study, we investigated the effects and interactions of early-life room temperature exposure and licking-like tactile stimulation on thyroid hormone receptor signaling in female rat pups. We focused on females in this study because there are possible sex differences in pup thermoregulation [25] and maternal care received [26]. In addition, the increase in estrogen during sexual differentiation in the male pup brain can interact with thyroid hormone for gene transcription [27]. We measured levels of circulating T3 and transcript abundance in the PVN of thyroid hormone receptors and the thyroid hormone-responsive genes *Dnmt3a* and *Oxt. We* hypothesized that early-life room temperature exposure and tactile stimulation would synergistically increase T3 levels as well as increase *Dnmt3a* and *Oxt* transcript abundance via changes in thyroid hormone receptor signaling. We predicted that the increase in *Oxt* transcript abundance would correspond to altered DNA methylation at the CHRE locus and differential thyroid hormone receptor binding. We also predicted that transcript abundance of other genes not directly regulated by thyroid hormone binding, including other DNA methyltransferases, *Crf* and *Avp*, would not show a similar pattern, and that *Crf* and *Avp* transcript abundance would be reduced with supplemental tactile stimulation.

## 2. Methods

### 2.1 Rat Breeding

Seven-week-old female (n = 28) and male (n = 16) Long-Evans rats were obtained from Charles River Laboratories (Kingston, NY, USA). They were housed in same-sex pairs on a 12:12 hour light-dark cycle (lights on at 7:00) with *ad libitum* access to standard chow diet and water. For breeding, one male was housed with two females for one week. Females were then housed separately and weighed weekly throughout pregnancy. All animal procedures were approved by the Local Animal Care Committee at the University of Toronto Scarborough and conformed to the guidelines of the Canadian Council on Animal Care.

Females were checked for parturition starting three weeks after breeding at 9:00 and 17:00. Postnatal day (PND) 0 was determined if the birth occurred between 9:00 and 17:00 or if pups were found at 9:00 but have not nursed yet. Pups found at 9:00 with a milk band were considered PND 1. At PND 1, litters were culled to four to six female pups and individually weighed. A total of 154 female rat pups were used for this study. One group of female rat pups was assessed for transcript abundance and DNA methylation and a separate group of female rat pups was assessed for triiodothyronine levels and chromatin immunoprecipitation enrichment.

### 2.2 Postnatal Manipulations

See Figure 1A for timeline of the experimental design between and within litters. From PND 2 to 7, nineteen whole litters were separated from their mother for approximately 25 minutes per day during the light phase (9:00 – 13:00) and placed in a small cage lined with corn cob bedding. Nine litters were placed in a cage warmed with a heating pad (set for 33-35° C; “nest temperature” condition) and ten litters were exposed to room temperature without extra heat (19-22° C; “repeated room temperature” condition). A total of 51 pups were in the nest temperature condition and 54 pups were in the repeated room temperature condition.

**Figure 1.**
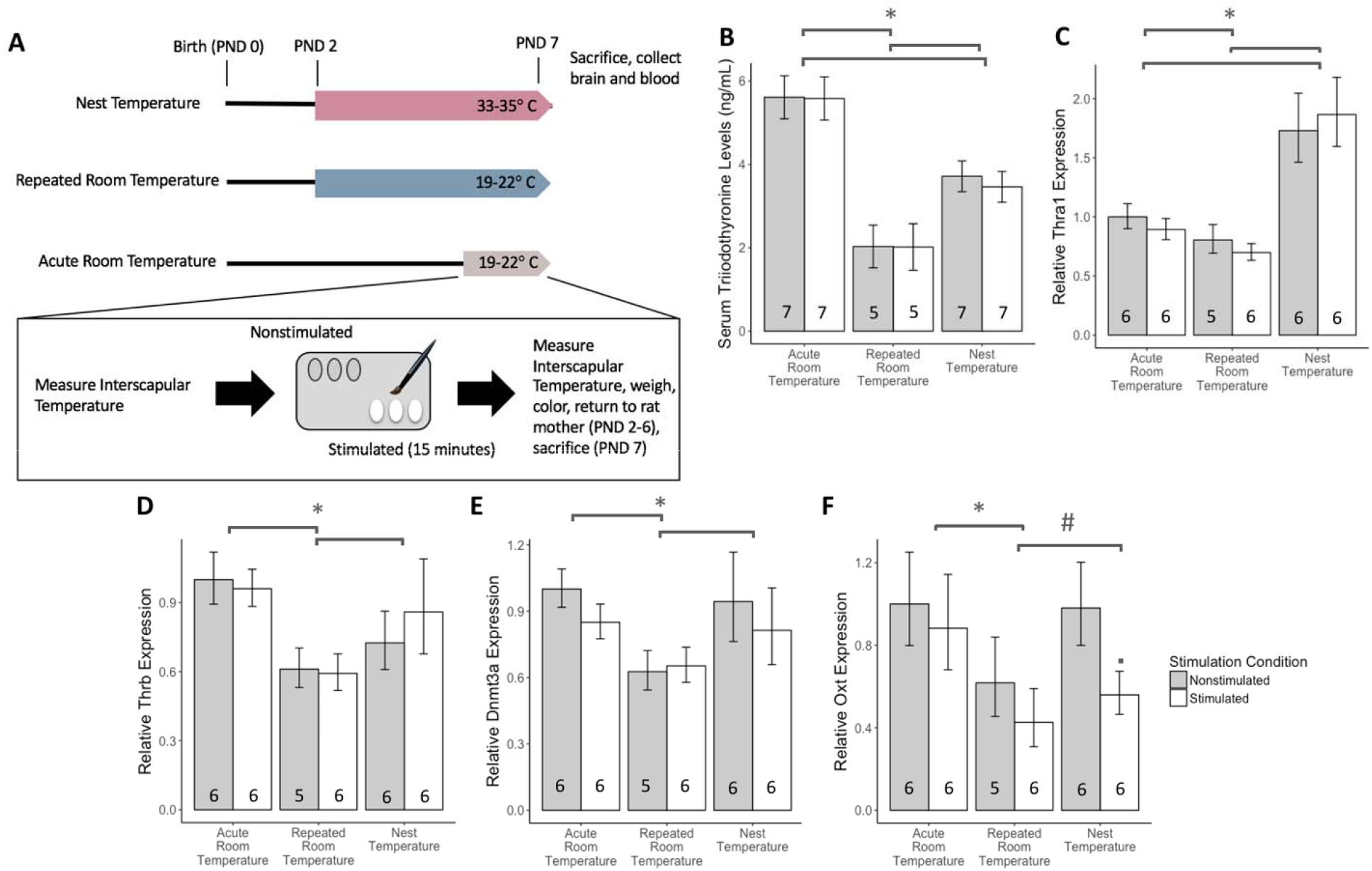
Repeated room temperature exposure decreased thyroid hormone activity with minimal effects of supplemental tactile stimulation. (A) Schematic diagram of the experimental design between litters and within litters of female rat pups. (B) Pups with repeated room temperature exposure had a significant decrease in circulating total triiodothyronine levels, (C) thyroid hormone receptor alpha 1 (*Thra1*) transcript abundance, (D) thyroid hormone receptor beta (*Thrb*) transcript abundance (E) DNA methyltransferase 3a (*Dnmt3a*) transcript abundance (F) and a marginal effect on oxytocin (*Oxt*) transcript abundance than rat pups exposed to nest temperature. Pups with acute exposure to room temperature had significant increases in circulating total triiodothyronine levels and thyroid-related transcript abundance than pups with repeated room temperature exposure. (F) Pups provided supplemental tactile stimulation had decreased *Oxt* transcript abundance in the nest temperature condition. The serum triiodothyronine barplot is displayed with mean +/- SEM. RT-qPCR barplots are displayed as fold changes relative to the pups handled once and nonstimulated +/- SEM. * p < 0.05, # p < 0.10 main effect of temperature condition; • p < 0.05 main effect of tactile stimulation condition.

Two to three female rat pups within a litter received supplemental tactile stimulation with a camel hair paintbrush (Craftsmart) for 15 minutes (“Stimulated” condition; n = 78) while the remaining pups were left undisturbed (“Nonstimulated” condition; n = 76). Within the nest temperature condition, 26 pups were in the stimulated condition and 25 pups were in the nonstimulated condition. Within the repeated room temperature condition, 27 pups were in the stimulated condition and 27 pups were in the nonstimulated condition. Pups received the supplemental tactile stimulation on the dorsal region of their body at a rate of approximately two strokes per second. The same pups received supplemental tactile stimulation between days. All pups were individually weighed daily and interscapular temperature was measured with an infrared thermometer (VWR) before and after the tactile stimulation period from PND 3 to 7. From PND 2 to 6, female rat pups were individually marked using odorless and tasteless food colouring (Club House, London, Ontario, Canada) to distinguish between siblings. This method to identify individual siblings has been implemented in previous work by our laboratory, and there have been no effects of the food colouring on maternal care [28].

To investigate whether the changes in the temperature exposure and tactile stimulation condition occurred over multiple separations in week-old pups, nine litters were separated once at PND 7 and two to three female rat pups within a litter received supplemental tactile stimulation. These litters were acutely exposed to room temperature without extra heat (“acute room temperature” condition). A total of 49 pups were in the acute room temperature condition; 25 pups were in the stimulated condition and 24 pups were in the nonstimulated condition. All pups were individually weighed and interscapular temperature was measured with an infrared thermometer (VWR) before and after the tactile stimulation period.

The interscapular temperature change (before the tactile stimulation period minus after the tactile stimulation period) was calculated for all groups at PND 7 to verify the room temperature exposure conditions had reduced pup temperature while the nest temperature exposure condition kept a stable pup temperature during the separation period.

At PND 7, all female rat pups were sacrificed following the tactile stimulation period and blood and brain were collected. One stimulated and one nonstimulated sibling were decapitated at a time. To examine if the time elapsed since the tactile stimulation period would affect the physiology of the pups, we noted the order pups were sacrificed for use as a control variable. Brains were flash frozen in isopentane and kept on dry ice. Whole blood samples were kept on ice to coagulate order at least 30 minutes before being centrifuged at 4000 x g at 4° C for 30 minutes. The serum was transferred to a new tube and the cell pellet was discarded. Both serum and brain were stored at −80° C. One to two pups per stimulation group per litter were used for all downstream molecular analyses.

### 2.3 Maternal Care Observations

From PND 2 to 6, each litter was video recorded for one hour two times during the light phase (13:00-14:00, 17:00-18:00) and three times during the dark phase (21:00-22:22, 1:00-2:00, 5:00-6:00). These videos were coded with Observer XT 10.5 (Noldus) for maternal behaviour by five coders with high inter-rater reliability (≥ 90%). Nursing, licking/grooming, nest-building, and other self-directed behaviours were scored every three minutes using an ethogram based on previous literature [29]. A total of 100 observations per day per mother were coded and each behaviour was represented as a percentage of the frequency of behaviour coded over total observations multiplied by 100. Total nursing was calculated as a sum of low crouch, high crouch, and supine nursing observations. Total licking was calculated as a sum of anogenital and body licking observations.

Two litters from the nest temperature condition had missing maternal care recordings at PND 2 and one litter from the repeated room temperature condition had missing maternal care recordings from PND 2-3 due to technical errors.

### 2.4 Serum Triiodothyronine Enzyme-Linked Immunosorbent Assay (ELISA)

The active thyroid hormone, triiodothyronine (T3), was measured in the pup serum (n = 5-7 per group) using an ELISA (MP Biomedicals Inc., USA) following the manufacturer’s instructions. For each pup sample, technical duplicates were measured when possible and 50 μl of serum per well was used. To keep all samples within the linear phase of the standard curve, the serum was diluted 1:1 with the “0” standard. Each plate was run and normalized with a control sample with a known concentration of T3 (Control Set I, Tri-level (for Steroid and Thyroid Hormones); MP Biomedicals Inc., USA). Concentration of T3 was determined using a 4-point logistic curve using an online software (https://elisaanalysis.com/app) and multiplied by two to account for the dilution factor.

### 2.5 Transcript Abundance with Quantitative Polymerase Chain Reaction (qPCR)

Postnatal day 7 pup brains (n = 5-6 per group) were cryosectioned with 50 μm slices using a Leica CM3050S cryostat. The paraventricular nucleus of the hypothalamus (PVN) (−1.40 to −2.00 mm Bregma) was microdissected using an atlas for the developing rat brain [30] and a supplementary atlas for the PND 7 rat brain [31]. RNA was extracted using a RNeasy Micro Kit (Qiagen) following the manufacturer’s instructions. Concentration and purity of RNA was assessed using a spectrophotometer (Nanodrop ND-2000C, Thermo Scientific). Up to 1 μg of RNA was converted to cDNA (Applied Biosystems High Capacity cDNA Conversion Kit) and was diluted to 5 ng/μl assuming 100% conversion efficiency.

Transcript abundance of candidate genes were assessed using StepOne Plus real-time PCR software with Fast SYBR Green PCR master mix (Applied Biosystems, Life Technologies, Carlsbad, CA, USA) using technical triplicates. Specifically, we analyzed arginine vasopressin (*Avp*), corticotropin releasing factor (*Crf*), DNA Methyltransferase 1 (*Dnmt1*), DNA Methyltransferase 3a (*Dnmt3a*), DNA Methyltransferase 3b (*Dnmt3b*), oxytocin (*Oxt*), thyroid hormone receptor α1 (*Thra1*), and thyroid hormone receptor β (*Thrb*). Each plate was run and corrected with one randomly assigned cDNA sample that was measured on all plates. All transcripts were normalized to the GEOmean of the Actin-β (*Actb*) and Ubiquitin C (*Ubc*) transcript and relative quantification was calculated by the ΔCT method. Supplementary Table S1 displays the custom-made primer sets created from Primer-BLAST software (National Center for Biotechnology Information) and previous literature [32,33].

### 2.6 DNA Methylation Analysis of the Oxytocin Promoter

DNA from six PVN from each supplemental tactile stimulation group in the repeated room temperature condition and nest temperature condition (total n = 24) was extracted using the Masterpure Complete DNA/RNA Extraction kit (Epicentre) and 300 ng of DNA was used for bisulfite conversion using the Epitect Fast Bisulfite Conversion kit (Qiagen) following the manufacturer’s instructions. Semi-nested PCR was performed with custom-made primers created from the Pyromark Q-CpG 1.0.9 software (Supplementary Table S1) and targeted one CpG site flanking the oxytocin composite hormone response element in the promoter region (chr3:123106520; rn6). The biotinylated amplicons were verified with gel electrophoresis and extracted with the MinElute Gel Extraction Kit (Qiagen). Pyrosequencing was done using a Pyromark Q106 ID pyrosequencer with technical triplicates and CpG methylation levels were calculated using the Pyromark Q-CpG 1.0.9 software.

### 2.7 Chromatin Immunoprecipitation (ChIP) with qPCR for DNA Methyltransferase 3a and Oxytocin

Two PVN from the same litter and supplemental tactile stimulation group were pooled (total n = 3 for repeated room temperature and n = 4 for nest temperature) for chromatin immunoprecipitation (ChIP) based on the protocol from Stefanelli and colleagues [34]. Two batches were done with 1-2 pooled PVN samples per temperature condition per batch.

Pooled PVN tissue samples were crosslinked with 1% formaldehyde (Sigma-Aldrich) for 10 minutes at 26° C. The samples were quenched with 1.25 M Glycine and left at room temperature for 5 minutes. The PVN tissue was pelleted using a centrifuge at 21,100 xg for 30 seconds before five washes with ice-cold PBS and a protease inhibitor cocktail (Roche) dissolved in PBS. The PVN tissue was homogenized with SDS lysis buffer (0.25 M Sucrose, 60 mM KCl, 15 mM NaCl, 10 mM MES (pH 6.5), 5 mM MgCl_2_, 0.5% Triton X-100) and centrifuged at 4,700 xg to pellet the cell nuclei. The SDS lysis buffer was removed and a salt buffer (50 mM NaCl, 10 mM PIPES (pH 6.8), 5 mM MgCl_2_, 1 mM CaCl_2_) was added prior to sonication on the lowest power setting (Output power: 3-4 Watts; 3x for 10 sec on, 30 sec off; Fisher Scientific Sonic Dismembrator Model 100). The samples were incubated with 150 units micrococcal nuclease (Cell Signaling) at 37° C for 10 minutes before quenching with 5 μl 0.5 M EDTA and placed on ice. 10% SDS was added to each sample before being centrifuged at 17,000 xg for 5 minutes, aliquoted, and diluted 4x with a ChIP dilution buffer. Each ChIP aliquot contained 20 μl of Millipore Protein G magnetic beads and 10 μg of thyroid hormone receptor α/β (Thermo-Fisher Scientific, cat no. MA1-215) or 2 μg of H3K27ac (Abcam, cat no. ab177178) antibody and incubated at 4° C overnight. The following day, the beads were pelleted using a magnetic separator and washed with ice-cold low-salt, high-salt, LiCl (Millipore), and Tris-EDTA buffers. Crosslinks were reversed for ChIP aliquots and input chromatin samples using 10 μg of proteinase K in Tris-EDTA buffer (with 1% SDS) at 65° C for at least 2 hours before purification using a PCR clean-up kit (BioBasic).

Primers for the *Oxt* composite hormone response element (CHRE) were created with Primer-BLAST software (Supplementary Table 1). Primers for the *Dnmt3a* 30.3 kbp and 49.3 kbp thyroid hormone response element (TRE) were created with Primer-BLAST software using the rat homologous sequences from previous literature using mouse ([16]; Supplementary Table S1). Enrichment was measured for each gene locus using qPCR for the ChIP DNA and input chromatin samples with technical triplicates. The enrichment for thyroid hormone receptors and H3K27ac were normalized and calculated as a relative percentage of input chromatin.

### 2.8 Statistical Analysis

All statistical analyses were performed using SPSS (IBM Corporation). To examine the effects of temperature condition on maternal care received within the first postnatal week, a repeated-measures 3 (Nest Temperature, Repeated Room Temperature, and Acute Room Temperature) x 5 (Postnatal days 2-6) linear mixed model was used to correct for missing datapoints and random factors. To examine the effects of room temperature exposure and tactile stimulation on female pup interscapular temperature change, serum T3 concentration, and transcript abundance, a 3 (Nest Temperature, Repeated Room Temperature and Acute Room Temperature) x 2 (Stimulated and Nonstimulated) general linear model was used to compare manipulation groups and their interactions. As there was a main effect of sacrifice order on *Thra1* and *Dnmt1* transcript abundance, a linear mixed model was used with order of sacrifice as a random factor. Significant effects of temperature condition were followed with a post-hoc test using Fisher’s Least Significant Differences. Significant effects of tactile stimulation or a significant temperature condition x tactile stimulation interaction were followed with a post-hoc test within each temperature condition. To examine the effects of room temperature exposure and tactile stimulation on DNA methylation of the *Oxt* promoter, a 2 (Nest Temperature and Repeated Room Temperature) x 2 (Stimulated and Nonstimulated) general linear model was used to compare manipulation groups and their interactions. To examine the effects of room temperature exposure on thyroid hormone receptor and H3K27ac enrichment on *Dnmt3a* and *Oxt*, a one-way (Nest Temperature and Repeated Room Temperature) linear mixed model was used with batch as a random factor. All effects were considered statistically significant at p ≤ 0.05 and marginally significant at p ≤ 0.10.

## 3. Results

### 3.1 Postnatal Day 7 Pup Characteristics and Maternal Care Received

There was a main effect of temperature exposure on interscapular temperature change at PND 7 (F _(2, 139)_ = 91.777, p < 0.001). Female rat pups in the nest temperature condition maintained a relatively stable interscapular temperature. In contrast, pups in both the acute and repeated room temperature conditions showed a significant reduction in interscapular temperature during the separation period. This reduction in temperature was greater for pups exposed to repeated room temperature compared to the pups in the acute room temperature condition (Supplementary Figure S2A).

There was no main effect of temperature exposure on total licking received (F _(2,25.383)_ = 1.498, p = 0.243; Supplementary Figure S2B) and total nursing received (F _(2,25.340)_ = 1.276, p = 0.296; Figure 2C) from the rat mother during the first postnatal week. However, total nursing significantly declined over the first postnatal week (F _(2,24.461)_ = 9.160, p < 0.001) and there was a significant temperature exposure x postnatal day interaction (F _(2,24.438)_ = 2.480, p = 0.040; Supplementary Figure S2C). Rat mothers with litters in the repeated room temperature condition and nest temperature condition provided significantly more nursing than mothers with litters in the acute room temperature condition at PND 4.

**Figure 2.**
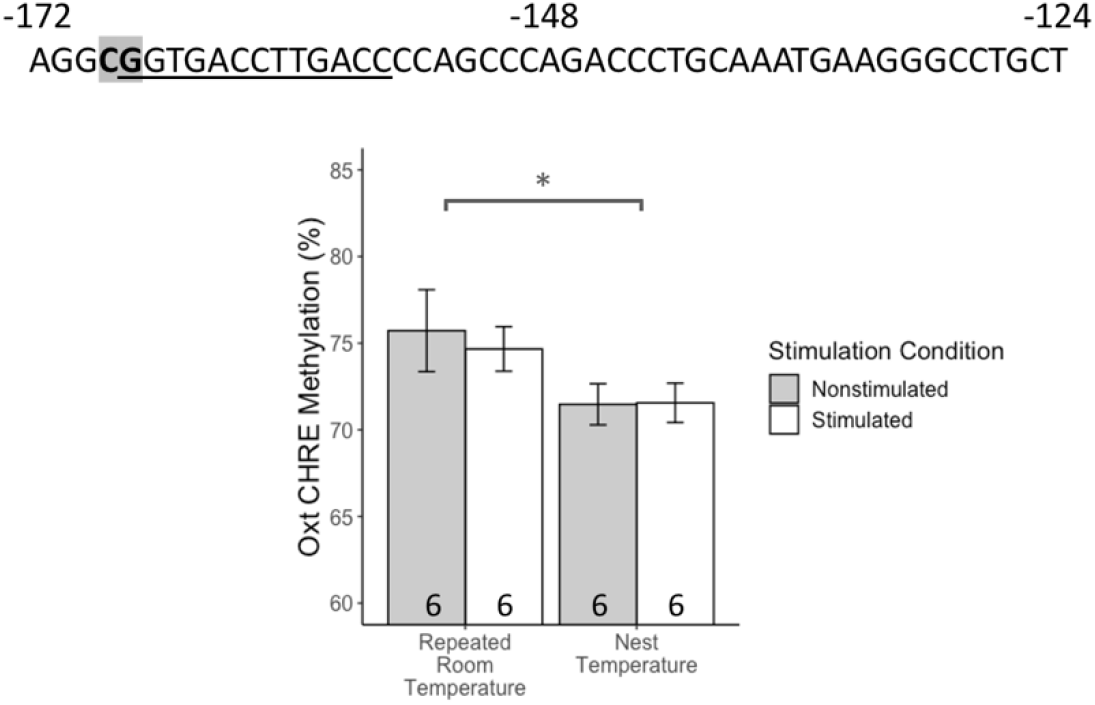
DNA methylation at the CpG site (highlighted) flanking the composite hormone response element in the oxytocin promoter (underlined) increased in female rat pups with repeated room temperature exposure. Barplot is displayed with mean +/- SEM. * p < 0.05 main effect of temperature condition.

### 3.2 Circulating Triiodothyronine Levels

There was a main effect of temperature exposure on T3 levels in the PND 7 rat pup serum (F _(2,42)_ = 35.141, p < 0.001; Figure 1B). Female rat pups in the repeated room temperature condition had significantly lower levels of T3 than pups in the nest temperature condition and acute room temperature condition (Post-Hoc p’s < 0.001). In addition, pups in the acute room temperature condition had higher levels of T3 than pups in the nest temperature condition (Post-hoc p < 0.001). There was no main effect of supplemental tactile stimulation and no interaction (p’s > 0.1) on serum T3 levels.

### 3.3 Transcript Abundance

Female rat pups with repeated room temperature exposure showed reduced thyroid hormone receptor signaling-related transcript abundance and rat pups with supplemental tactile stimulation showed reduced *Oxt* transcript abundance. There were main effects of temperature exposure on *Thra1* (F _(2, 28.755)_ = 113.774, p < 0.001; Figure 1C), *ThrB* (F _(2, 29)_ = 13.233, p < 0.001; Figure 1D), *Dnmt3a* (F _(2, 29)_ = 8.481, p =0.001; Figure 1E), and *Oxt* (F _(2, 29)_ = 3.794, p = 0.034; Figure 1F) transcript abundance. Specifically, pups in the repeated room temperature condition had a significant reduction of *Thra1* (Post-hoc p < 0.001), *Thrb* (Post-hoc p < 0.001), *Dnmt3a* (Post-hoc p = 0.001), and *Oxt* (Post-hoc p = 0.008) transcript abundance compared to pups in the acute room temperature condition. Pups in the repeated room temperature condition also had a significant reduction of *Thra1* (Post-hoc p < 0.001), *Thrb* (Post-hoc p = 0.011), *Dnmt3a* (Post-hoc p = 0.003), and a marginal reduction in *Oxt* (Post-hoc p = 0.092) transcript abundance compared to pups in the nest temperature condition. In addition, pups in the acute room temperature condition had a significant reduction of *Thra1* (Post-hoc p < 0.001) and a significant increase of *Thrb* (Post-hoc p = 0.018) transcript abundance compared to the nest temperature condition.

There was a marginal effect of supplemental tactile stimulation on *Oxt* transcript abundance (F _(1, 29)_ = 3.890, p = 0.058). Pups with supplemental tactile stimulation had a decrease in *Oxt* transcript abundance if they were in the nest temperature condition (F _(1,10)_ = 7.011, p = 0.024; Figure 1F). There were no main effects of temperature or tactile stimulation on *Avp* (Supplementary Figure S3A) and *Crf* (Supplementary Figure S3B) transcript abundance and no interactions in any of the genes measured (p’s > 0.1).

There was also a main effect of temperature exposure on *Dnmt1* transcript abundance (F _(2, 28.757)_ = 9.233, p = 0.001; Supplementary Figure S3C). Female rat pups with repeated room temperature exposure had a significant reduction in *Dnmt1* transcript abundance relative to pups in the acute room temperature condition (Post-hoc p = 0.015) and pups in the nest temperature condition (Post-hoc p < 0.001). There was a marginal reduction in *Dnmt1* among pups in the acute room temperature condition compared to pups in the nest temperature condition (Post-hoc p = 0.087). There were no main effects of temperature or tactile stimulation on *Dnmt3b* transcript abundance (Supplementary Figure S3D).

### 3.4 Oxytocin DNA Methylation Levels at the Composite Hormone Response Element

To investigate the long-term changes of room temperature exposure and supplemental tactile stimulation, we restricted the analysis on DNA methylation levels to the rat pups with repeated early-life separations. Female rat pups with repeated room temperature exposure had increased DNA methylation levels flanking the composite hormone response element (CHRE) in the *Oxt* promoter compared to pups in the nest temperature condition (F _(1,20)_ = 5.256, p = 0.033; Figure 2). There was no main effect of supplemental tactile stimulation and no interaction (p’s > 0.1) on DNA methylation levels.

### 3.5 Chromatin Immunoprecipitation Enrichment of Thyroid Hormone Receptor and H3K27ac

The *Oxt* CHRE in the promoter region and two thyroid response elements (TRE) within the *Dnmt3a* gene (Figure 3A) were analyzed for thyroid hormone receptor binding and H3K27ac levels. Female rat pups in the repeated room temperature condition had a significant increase in thyroid hormone receptor enrichment at the *Oxt* CHRE relative to pups in the nest temperature condition (F _(1,4.015)_ = 17.811, p = 0.013; Figure 3B). There were no effects effects of repeated temperature at the *Dnmt3a* +30.3 kbp TRE (F _(1,4.046)_ = 0.022, p = 0.889) or +49.3 kbp TRE (F _(1,4.031)_ = 0.234, p = 0.654). Likewise, there were no effects of temperature on H3K27ac enrichment in any gene loci tested (p’s > 0.1; Figure 3C).

**Figure 3.**
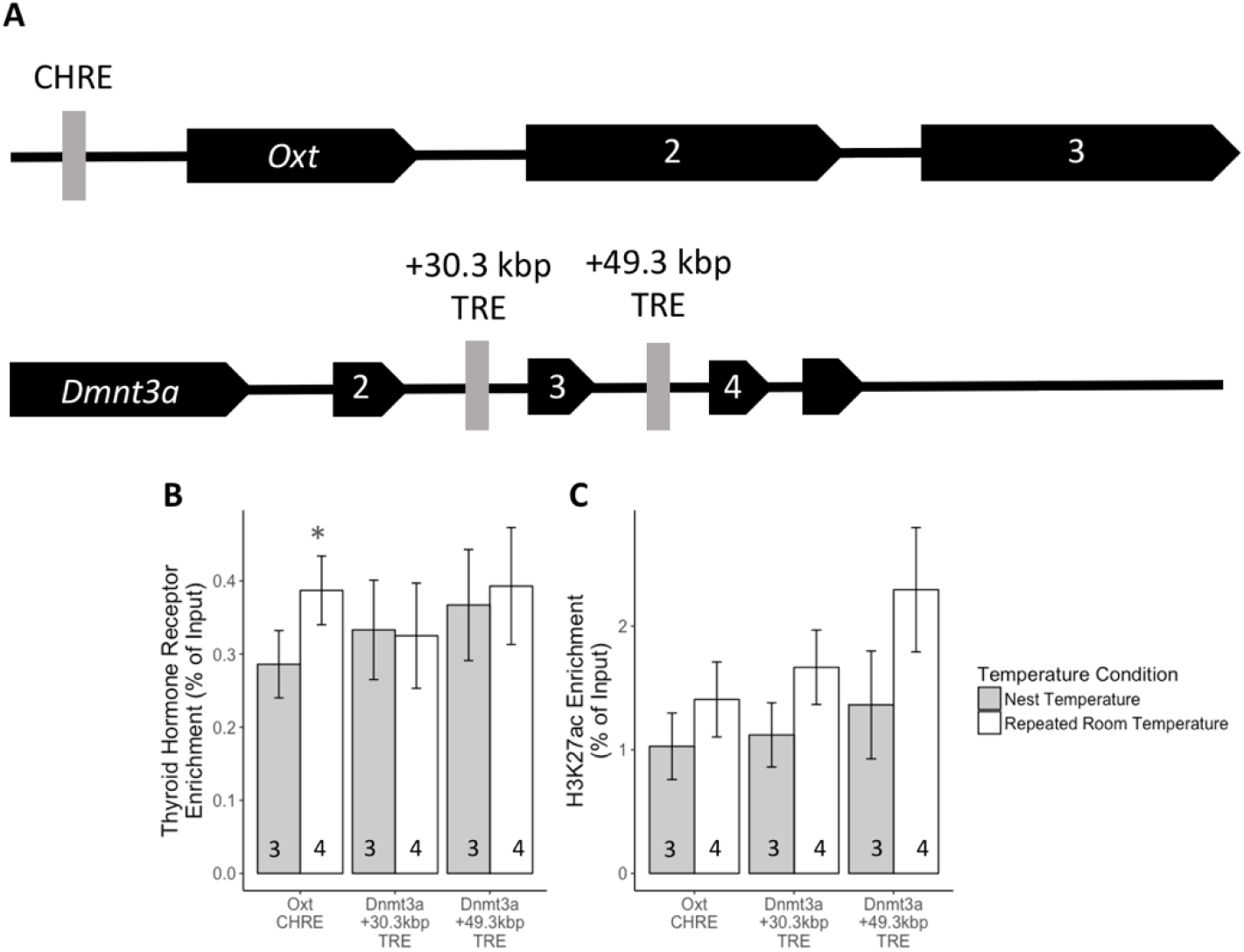
Early-life temperature exposure affected thyroid hormone receptor binding at the oxytocin composite hormone response element (CHRE) and did not change histone levels of H3K27ac in the paraventricular nucleus. (A) Schematic diagram of regulatory gene loci tested for *Oxt* and *Dnmt3a*. (B) Female rat pups with repeated room temperature exposure had significant increases in thyroid hormone receptor enrichment at the *Oxt* CHRE but not at the *Dnmt3a* thyroid hormone response elements (TRE) with (C) no significant changes in H3K27ac enrichment in any gene loci. Barplot is displayed with mean +/- SEM. * p < 0.05 main effect of temperature condition.

## 4. Discussion

In this study, we investigated how two factors rat pups commonly experience in the early-life maternal environment, ambient (room) temperature exposure and licking-like tactile stimulation, would affect neurodevelopment by changes in thyroid hormone receptor signaling at the neonate stage. This is the first study to our knowledge to investigate the contribution of early-life room temperature exposure on epigenetic modifications in the brains of rat pups. We found that female rat pups subjected to repeated room temperature exposure in early-life showed a reduction in several measures of thyroid hormone receptor signaling relative to pups with early-life nest temperature exposure, including circulating triiodothyronine and transcript abundance in the PVN of thyroid hormone receptors and the thyroid hormone-responsive genes *Dnmt3a* and *Oxt*. These effects were associated with increased DNA methylation and thyroid receptor binding at the composite hormone response element in the *Oxt* promoter in room temperature-exposed pups. Female rat pups with acute room temperature exposure showed the highest levels of T3 and an increase in measures of thyroid hormone receptor signaling tested relative to rat pups with repeated room temperature exposure. There was no effect of supplemental tactile stimulation on most of the thyroid hormone receptor signaling measures and minor effects on transcript abundance of *Oxt* in the PVN. These findings indicate that early-life room temperature exposure, a proxy for reduced maternal contact, may influence offspring phenotype via changes in thyroid hormone receptor signaling and downstream DNA methylation modifications. In addition, our results suggest that the changes in *Dnmt3a* transcript levels and DNA methylation at the *Oxt* CHRE occur as a consequence of alterations in thyroid hormone receptor signaling.

### 4.1 Effects on Thyroid Hormone Receptor Signaling

We predicted that room temperature exposure would increase circulating triiodothyronine levels in order to activate thermogenesis. However, we found decreased circulating triiodothyronine in response to repeated room temperature exposure, as well as decreased transcript abundance of thyroid hormone receptors and the thyroid hormone-responsive genes *Dnmt3a* and *Oxt*, compared to pups with nest temperature exposure. Though the decrease in *Oxt* is a nonsignificant trend, it followed a general pattern of repression of thyroid hormone receptor signaling with an increase in DNA methylation in the *Oxt* promoter region. Female rat pups exposed to acute room temperature showed the predicted increase in these measures, demonstrating that a suppression in thyroid hormone receptor signaling occurs with repeated exposures to room temperature. We also observed minimal alterations in maternal care received between the different temperature exposure groups; therefore, the data suggest that these changes in the rat pups are more likely due to the temperature manipulations directly than to indirect changes in maternal care received.

Cold acclimatization in adult rats can decrease levels of thyroid hormone released in response to a cold stressor [35,36], possibly as thyroid hormone is no longer required to activate thermogenesis [37]. Interestingly, rat pups exposed to repeated room temperature also showed a more pronounced decrease in interscapular temperature, the main site for thermogenesis, than rat pups with an acute exposure to room temperature during the separation period at PND 7. It is unknown if these changes in thyroid hormone physiology persist into adulthood and if they would affect other phenotypes. However, studies in mice show a potential link between nest quality, with lower quality nests associated with greater exposure of the pups to the ambient temperature, and increased metabolic rate, thermogenesis, and thyroxine levels at adulthood [38,39]. Overall, these findings suggest that the effects of repeated room temperature exposure in early life may reflect a physiological adaptation to cold stressors in the rat pup.

We also found that repeated room temperature exposure was associated with decreased transcript abundance of *Dnmt1*, which has not been previously shown to be responsive to thyroid hormone. However, one study has shown *Dnmt3a* and *Dnmt1* can cooperatively add *de novo* methyl groups to both strands of DNA [40]. Incubation of a DNA fragment with *Dnmt3a* before *Dnmt1* will stimulate DNA methylation modifications while the inverse does not, suggesting that this relationship is mainly driven by changes in *Dnmt3a* [40], though follow-up studies have not been done to our knowledge. Therefore, it is possible that the changes in *Dnmt1* transcript abundance in our study were a downstream effect of the changes in *Dnmt3a* transcript abundance.

We did not find effects of supplemental tactile stimulation on most of the thyroid hormone receptor signaling measures tested, which contrast from the study results by Hellstrom and colleagues [14]; however, their study used male rat pups as subjects with one instance of supplemental tactile stimulation for five minutes. It is possible that the effects of tactile stimulation on T3 levels are transient and the deiodinase activity decreased over prolonged periods of tactile stimulation in our study. It is also possible that female rat pups respond differently to tactile stimulation than male pups, given that males and females can respond differently to natural variations in maternal care received [41].

We found that supplemental tactile stimulation was associated with a trend in decreased *Oxt* transcript abundance, though this did not correspond to changes in DNA methylation in the CHRE. We also find similar but nonsignificant decreases in *Crf* and *Avp* transcript abundance. There is some evidence that tactile stimulation can increase oxytocinergic neuron activity [42] and that variation in maternal care can be transmitted across generations through changes in the oxytocinergic system in female offspring [43,44]. However, other studies have shown that augmented maternal care and brief early-life separations *decrease* oxytocin transcript abundance and oxytocin-positive neurons in the hypothalamus [45,46]. In addition, one study showed that the proximal thermotactile contact with the rat mother but not maternal licking increased oxytocin neuropeptide concentrations in the hypothalamus [47]. Therefore, the relationship between maternal care received and pup oxytocin appears to be complex, and temperature exposure may be one confounding factor in these studies.

### 4.2 Effects on DNA methylation and the Role of Thyroid Hormone Receptor

We hypothesized that changes in transcript abundance of *Oxt* and *Dnmt3a* would be mediated by differences in DNA methylation (in oxytocin) and thyroid hormone receptor binding. We found that a decrease in *Oxt* transcript abundance in the repeated room temperature exposure condition corresponded to increased levels of DNA methylation and thyroid hormone receptor binding at the CHRE in the promoter region of oxytocin. There is some evidence that DNA methylation changes associated with variations in maternal care received occur as a consequence of differential transcription factor binding [6]. One possible explanation of our findings is that there is a repressive effect of unliganded thyroid hormone receptor on *Oxt* transcription. If unliganded by T3, thyroid hormone receptor can recruit repressors, including silencing mediator of thyroid and retinoic receptors (SMRT) and nuclear receptor corepressor 1 (NCOR1), and activate the histone deacetylase HDAC3 to silence gene transcription [48,49]. HDAC3 appears to mainly affect H3K9ac levels [50], which may explain the nonsignificant differences in H3K27ac observed in our study. However, no studies to our knowledge have found HDAC3 activity preceding DNA methyltransferase binding or DNA methylation modifications. These findings imply that higher DNA methylation at the oxytocin CHRE may occur as a consequence of unliganded thyroid hormone receptor binding, but future work is needed to elucidate this hypothesis more directly.

While repeated room temperature exposure decreased *Dnmt3a* transcript abundance, this was not associated with differences in thyroid hormone receptor binding in the regulatory TRE regions tested. We tested these two sites because of previous evidence of differential binding of thyroid hormone receptors in regulating *Dnmt3a* transcript in the developing mouse brain [16]. However, other TREs exist in the *Dnmt3a* gene and their responsiveness to thyroid hormone can be species specific [16]. Future studies are needed to verify which *Dnmt3a* TREs would be most relevant for the transcription of *Dnmt3a* in the neonatal rat brain.

## 5. Conclusions

Overall, our findings indicate that early-life room temperature exposure may affect DNA methylation indirectly by changes in DNA methyltransferases and directly by modifications at specific gene loci (oxytocin CHRE) as a consequence of differences in thyroid hormone receptor signaling. There is accumulating evidence that the epigenome is dynamic and responsive to environmental exposures, but studies that have investigated the underlying mechanisms between environmental exposures and DNA methylation are limited. In addition, there is evidence that DNA methyltransferase recruitment is targeted based on transcriptional activity and transcription factor binding at specific gene loci [51,52]. Other studies have proposed that the effects of other steroid hormones are mediated by the their effects on DNA methylation modifications, either through differential steroid receptor binding or DNA methyltransferase activity [53,54], typically in light of transcriptional activation. Our findings support the hypothesis that steroid hormones can affect the epigenome through both steroid receptor binding and DNA methyltransferase activity as well as induce transcriptional repression.

