## Supplemental Table and Figures for "Early-life Temperature Exposure Affects Thyroid Hormone Receptor Signaling and Epigenetic Regulation of the Paraventricular Nucleus in Female Rat Pups"

**Table S1.** List of primers and sequences used for qPCR, bisulfite sequencing, and ChIP.

| Gene | Forward Primer Sequence (5' to 3') | Reverse Primer Sequence (5' to 3') | Reference |
| --- | --- | --- | --- |
| **qPCR** | | | |
| Actb | ACCCGCGAGTACAACCTTCTTG | TATCGTCATCCATGGCGAACTGG |  |
| Avp | CATCCGACATGGAGCTGAGAC | CCACGCAGCTCTCATCGCT |  |
| Crf | TTCCTGTTGCTGTGAGCTTG | TCACCTTCCACCTTCTGAGG | [32] |
| Dnmt1 | ACCTACCACGCCGACAT | AGGTCCTCTCCGTACTCCA |  |
| Dnmt3a | ACGCCAAAGAAGTGTCTGCT | CTTTGCCCTGCTTTATGGAG |  |
| Dnmt3b | GATGTGACACCTAAGAGCAGCAGTAC | CAAACTCCTTGTCATCCTGATACTCA |  |
| Oxt | TGCGCAAGTGTCTTCCCTGCG | AGCCATCCGGGCTACAGCAGA | [33] |
| Thra1 | AAGCCAAGCAAGGTGGAGTGTG | ATCTGGTGACCTGGCACTGTTC |  |
| Thrb | AGCTCTGGCATTCCCTTATTCA | ATCCGTGGTTTCCCTCTCCT |  |
| Ubc | CACCAAGAAGGTCAAACAGGAA | AAGACACCTCCCCATCAAACC |  |
| **Bisulfite Pyrosequencing** | | | |
| Oxt (Outer) | AGTTTTATTTTGAGGTATTGGATTTTATG | CACACTATTTAAAAACAAACCCTTCATT |  |
| Oxt (Semi-Nested) |  | Biotin-ACTATTTAAAAACAAACCCTTCATTTACA |  |
| Oxt (Sequencing) | TTGAGTTTTAGGTTATTAGTTG | |  |
| **ChIP** | | | |
| Dnmt3a  (+30.3 kbp TRE) | GGAAGTCAGATGAGGTCACGG | CCAGAGCCAGAGCAGTTACTA |  |
| Dnmt3a  (+49.3 kbp TRE) | CACCAACTTCCTCCAGGGTTA | GCCCTTGACGGGTTATTCTT |  |
| Oxt (CHRE) | CTCCAGGTCATTAGCTGAGGC | TGCATGACTGGTCACAGCAG |  |

***Actb***: Actin beta, ***Avp***: Arginine vasopressin, ***Crf***: Corticotropin releasing factor, ***Dnmt1***: DNA methyltransferase 1, ***Dnmt3a***: DNA methyltransferase 3a, ***Dnmt3b***: DNA methyltransferase 3b, ***Oxt***: oxytocin, ***Thra1***: Thyroid hormone receptor alpha-1, ***Thrb***: Thyroid hormone receptor beta, ***Ubc***: Ubiquitin C, **CHRE**: Composite hormone response element, **TRE**: Thyroid hormone response element

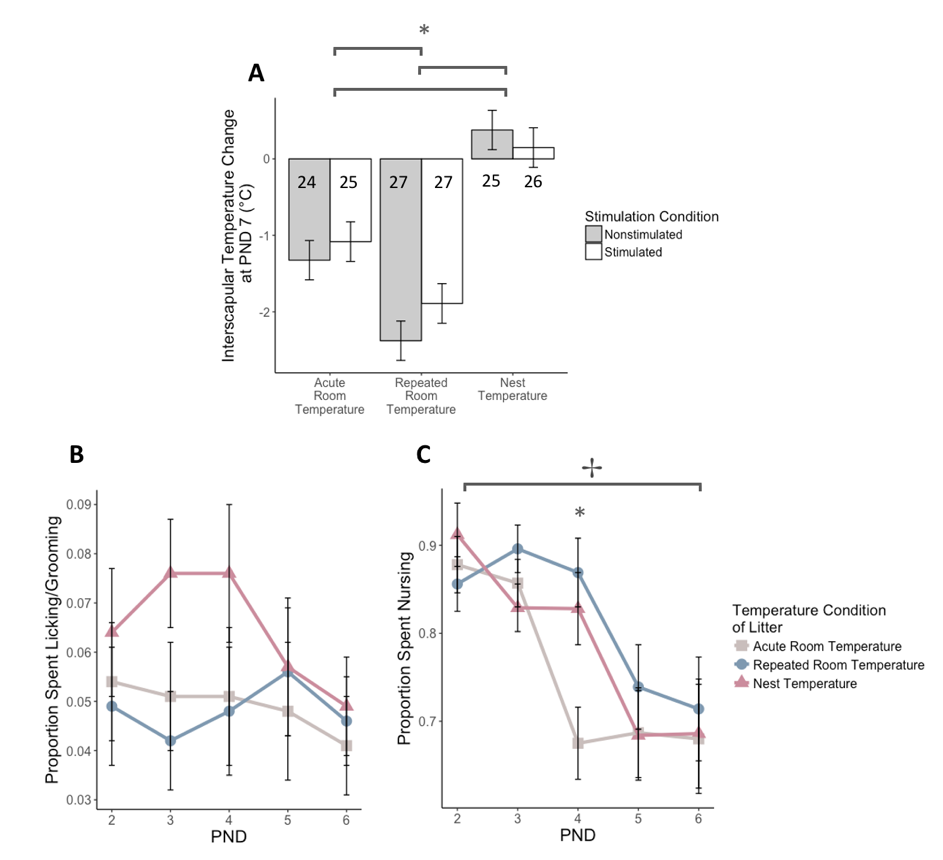

**Figure S1.** Early-life temperature exposure significantly affected temperature change and weight in week-old female pups with minor differences in maternal care received. (A) Female rat pups exposed to repeated room temperature had a significantly more pronounced reduction in body temperature compared to pups exposed to acute room temperature. Both groups had a more pronounced reduction in body temperature than pups exposed to nest temperature. (B) Litters in the different temperature conditions received similar amounts of maternal licking/grooming during the first postnatal week. (C) Nursing decreased during the first postnatal week and the litters in the acute room temperature condition receiving less nursing at postnatal day (PND) 4 than litters exposed to repeated room temperature or nest temperature. Barplots are shown with mean +/- SEM and line graphs are shown with estimated marginal means +/- SEM. * p < 0.05 main effect of temperature condition, ✢ p < 0.05 main effect of postnatal day.

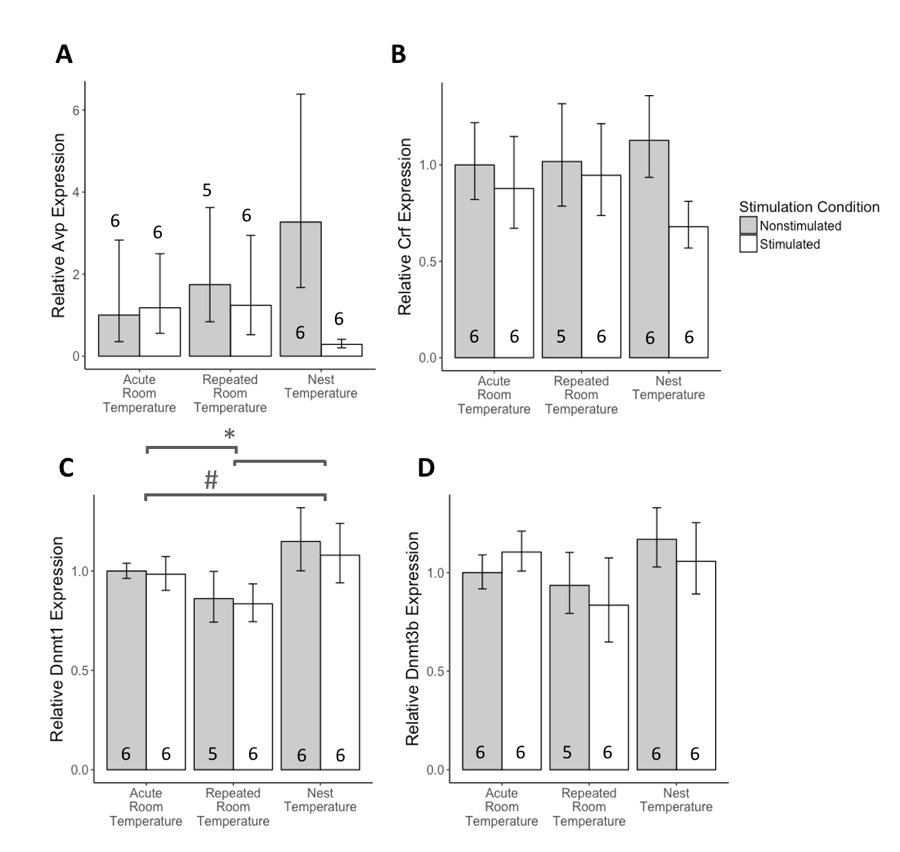

**Figure S2.** Early-life temperature exposure affected DNA methyltransferase 1 (*Dnmt1*) transcript abundance and neither early-life temperature exposure nor supplemental tactile stimulation affected DNA methyltransferase 3b (*Dnmt3b*) and other stress-related neuropeptides in the paraventricular nucleus. (A) Female rat pups with repeated room temperature exposure had a significant decrease in *Dnmt1* transcript abundance than rat pups acutely exposed to room temperature and rat pups exposed to nest temperature. There are no main effects or interaction of temperature exposure and supplemental tactile stimulation on (B) *Dnmt3b*, (C) arginine vasopressin (*Avp*) or (D) corticotropin releasing factor (*Crf*) transcript abundance. RT-qPCR barplots are displayed as fold changes relative to the pups handled once and nonstimulated +/- SEM. * p < 0.05, # p < 0.10 main effect of temperature condition.
